# The Interplay of Acetylation and Ubiquitination Controls PRMT1 Homeostasis

**DOI:** 10.1101/2024.06.18.599616

**Authors:** Mohd. Altaf Najar, Jenna N. Beyer, Callie E. W. Crawford, George M. Burslem

## Abstract

PRMT1 plays many important roles in both normal and disease biology, thus understanding it’s regulation is crucial. Herein, we report the role of p300-mediated acetylation at K228 in triggering PRMT1 degradation through FBXL17-mediated ubiquitination. Utilizing mass-spectrometry, cellular biochemistry, and genetic code-expansion technologies, we elucidate a crucial mechanism independent of PRMT1 transcript levels. These results underscore the significance of acetylation in governing protein stability and expand our understanding of PRMT1 homeostasis. By detailing the molecular interplay between acetylation and ubiquitination involved in PRMT1 degradation, this work contributes to broader efforts in deciphering post-translational mechanisms that influence protein homeostasis.

## Introduction

Protein arginine methyltransferase 1 (PRMT1) is the prototypical member of the PRMT enzyme family, which catalyze the transfer of a methyl group from *S*-adenosyl methionine (SAM) to arginine sidechains (Beyer et al., 2021; Greenblatt et al., 2016). PRMT1 is responsible for the majority of asymmetric dimethylation of arginine residues in mammalian cells (Pawlak et al., 2000; Tang et al., 2000). Most notably, PRMT1 methylates Arginine 3 on histone H4 (H4R3me2a) which recruits and activates lysine acetyltransferases CBP/p300 to acetylate neighboring histone tails, resulting in chromatin relaxation and thus enhanced transcription (An et al., 2004; Huang et al., 2005; Li et al., 2010). PRMT1 also has many non-histone targets involved in transcription, translation, RNA processing, DNA damage response, and signal transduction (Jiang et al., 2023a; Sudhakar et al., 2024; Thiebaut et al., 2021). PRMT1’s central role in many cellular processes means that mis-regulation has diverse consequences and as such, PRMT1 has been implicated in a variety of diseases (Wu et al., 2021; Yang and Bedford, 2013).

PRMT1 has largely been characterized as a critical driver of breast cancer through its roles in transcription (Gao et al., 2016; Li et al., 2021) and methylation of estrogen and progesterone receptors (Le Romancer et al., 2008; Malbeteau et al., 2020) as well as regulation of epidermal growth factor receptor and Wnt signaling pathways (Suresh et al., 2022). Dysregulation of these processes has also been shown to drive progression in other cancers, such as colorectal cancer (Liu et al., 2024; Zhang et al., 2023b), melanoma (Tao et al., 2024), hepatocellular carcinoma (Jiang et al., 2023b), prostate cancer (Tang et al., 2022), pancreatic cancer (Nguyen et al., 2024), leukemia (Cheung et al., 2007; Shia et al., 2012), and non-small cell lung cancer (Avasarala et al., 2015). More recently, PRMT1 has been linked to cancer metabolism through promoting glycolysis in hepatocellular carcinoma and colorectal cancer (Li et al., 2023; Liu et al., 2024), as well as fatty acid metabolism in hepatocellular carcinoma and breast cancer (Yamamoto et al., 2024; Yan et al., 2024). PRMT1 has also been shown to play an important roles beyond oncology, including in cardiovascular disease (Couto e Silva et al., 2020), diabetes (Iwasaki, 2024) and neurodegenerative diseases (Angelopoulou et al., 2023) such as amyotrophic lateral sclerosis (Aikio et al., 2021). Many of the above studies have shown that knockdown or knockout of PRMT1 *in vitro* and *in vivo* has significant effect on disease phenotypes. However, inhibition of PRMT1 has failed to fully recapitulate these phenotypes (Cheung et al., 2016; Shia et al., 2012). Further, clinical trials of PRMT1 inhibitors have failed due to lack of response and toxicity (El-Khoueiry et al., 2023). Together, this suggests that the roles of the protein in disease may go beyond its enzymatic activity (Hsu et al., 2017), motivating us to understand the mechanisms of PRMT1 homeostasis as a potential mechanism to modulate expression levels.

The canonical mechanism for protein degradation in eukaryotic cells involves the deposition of a K48-linked ubiquitin chain onto the surface of a target protein via an E1-E2-E3 enzymatic cascade, resulting in 26S proteasomal degradation. While E1 and E2 enzymes work to activate and conjugate ubiquitin, E3 ligases enact the final step of ubiquitin attachment by directly binding the target protein and facilitating ubiquitin transfer onto an exposed lysine residue. E3 ligases are therefore largely responsible for conferring substrate specificity to degradation pathways and are often controlled by post-translational modifications (PTMs) that constitute or hinder E3 ligase binding to the target protein (Lee et al., 2023). Here, we reveal that PRMT1 degradation can be controlled by an acetylation event on Lysine 228, triggering recognition and subsequent ubiquitination by E3 ligase SCF-FBXL17, a member of the F-Box Cul1 family (Willems et al., 2004). We validate this finding using an innovative application of genetic code expansion in mammalian cells, site specifically incorporating acetyl-lysine at sites in PRMT1.

## Materials and Methods

### Materials

Trypsin/Lys-C Mix, Mass Spec Grade (V5072) was purchased from Promega (Madison, Wisconsin, USA). Pierce™ Quantitative Colorimetric Peptide Assay (23275) and Pierce™ Rapid Gold BCA Protein Assay Kit (A53227) were ordered from ThermoFisher Scientific (Waltham, Massachusetts, USA). ZipTip with C18 resin (ZTC18S096) was ordered from Millipore (Burlington, Massachusetts, USA). Antibodies anti-DYKDDDDK Tag (9a3) (8146s), anti-acetyl-Lysine (9441s), anti-PRMT1 (a33) (2449S), anti-K48 linkage specific (d9d5) (8081t), anti-K63 linkage specific (d7a11) (5621t), anti-p300 (e8s2v) (57625s), anti-PRMT1 (E5A8F) (79780s), anti-DYKDDDDK Tag (d6w5b) (86861s), anti-acetyl-Histone H3 (k27) (d5e4) xp(r) (8173s), anti-Histone H3 (d1h2) xp(r) (4499t), anti-Myc-tag (71d10) (14038s), anti-HA-tag (c29f4 and 3724s), anti-HA-tag HRP conjugated (c29f4 and 14031s), anti-V5-tag (d3h8q) (1320s), and Rabbit mAb (Magnetic Bead Conjugate) (31628S) were obtained from Cell Signaling Technology (Danvers, Massachusetts, USA). Anti-β actin–HRP conjugated antibody was purchased from Sigma-Aldrich (St. Louis, Missouri, USA). Anti-FBXL17 (ab111683) was ordered from Abcam (Boston, Massachusetts, USA). Dulbecco’s Modified Eagle Medium (DMEM) with high glucose, fetal bovine serum (FBS), antibiotic antimitotic (PenStrep) solution and trypsin were purchased from Gibco (ThermoFisher Scientific, USA). A-485 (24119) was ordered from Cayman Chemicals (Ann Arbor, Michigan, USA). Deubiquitinase inhibitors PR-169 (A8212) were ordered from APExBIO (Boston, Massachusetts, USA). Proteasome inhibitor MG-132 (474787-10MG) was ordered from Sigma-Aldrich. Deacetylation Inhibition Cocktail (HY-K0030) was ordered from Medchemexpress (Monmouth Junction, New Jersey, USA). Cycloheximide was ordered both from Sigma-Aldrich (01810-1G) and Cell Signaling Technology (2112S). ANTI-FLAG® M2 affinity gel (A2220-10ML), Anti-C-Myc Magnetic Beads (SAE0201-1ML), and FLAG® peptide (F3290-4MG) was purchased from Sigma-Aldrich. Pierce™ Anti-HA magnetic beads (88837) were purchased from ThermoFisher Scientific. Solvents and consumables for LC-MS/MS were ordered from ThermoFisher Scientific. All other consumables used in the study were from Sigma-Aldrich, United States. PolyJet (SL100688) was ordered from SignaGen laboratories (Houston, Texas, USA). Cell culture plasticware ordered from Fisherbrand, Thermo Scientific, Millipore, and Falcon.

## EXPERIMENTAL MODEL AND SUBJECT DETAILS

### Cell culture

HEK293T, HEK293T FLAG-PRMT1, HEK293T FBXL17 KO and HEK293T acetyl-lysine incorporation cells were maintained in DMEM supplemented with 10% FBS, 100 U of penicillin, and 100 mg/mL streptomycin. All cells were cultured in a humidified CO_2_ incubator, 5% CO2 in a temperature at 37°C.

### Generation of FBXL17 knockout cell lines

HEK293T cells were transfected using PolyJet (SignaGen) with human FBXL17CRISPR-Casp9-Guide_LentiCRISPR-V2.0 Plasmids with both Cas9 and FBXL17 guide were ordered from Genscript (Catalogue SC1678). At 48 h after transfection, cells were selected in puromycin (1 µg/mL)-containing medium for 3 days. The resulting cells were subjected to clonal isolation by the single cell dilution method. The knockout of *FBXL17* was validated by immunoblot analysis.

### FLAG-PRMT1-Stable Expressing Cells

HEK Flp-In-293 cells (Invitrogen) were maintained in DMEM supplemented with 10% (*v/v*) heat-inactivated FBS (Gibco ThermoFisher Scientific, United States) and 1%b Pen-Strep (Gibco) in 5% (*v*/*v*) CO_2_. The HEK Flp-In-293 cells were transfected with the pcDNA5/FRT/TO - PRMT1 expression vectors and the Flp recombinase expression plasmid pOG44 using PolyJet (SignaGen). Colonies resistant to 100 µg/mL hygromycin B were picked and sub-cultured. The selected cells were maintained in DMEM supplemented with 10% (*v*/*v*) heat inactivated FBS and in 5% (*v*/*v*) CO_2_. 1 µg/mL tetracycline was used to induce the expression of FLAG-tagged PRMT1. Expression of FLAG-PRMT1 was validated by immunoblotting.

### Genetic code expansion cell line for acetyl-lysine incorporation

To generate a cell line for acetyl-lysine incorporation, we followed a previously established protocol (Elsässer et al., 2016). Briefly, ∼570,000 HEK293T cells were co-transfected with 500 ng of plasmid containing SuperPiggyBac transposase and 2.5 µg of a plasmid, referred to as E312, containing an *M. mazei* amino-acyl tRNA synthetase for AcK and 5 copies of the orthogonal *M. mazei* tRNA. PolyJet (SignaGen) was used for the transfection. Cells were left to recover for 2 days, before being split 1:5. The following day, 0.5 µg/mL of Puromycin (Sigma-Aldrich) was added to the media, and selection was continued for a total of 5 days. Cells were then returned to complete DMEM and expanded as a polyclonal pool.

## METHOD DETAILS

### Immunoblots and immunoprecipitation

Cells were lysed in lysis buffer (50 mM HEPES pH 7.5, 100 mM KCl, 2 mM EDTA, 0.3% NP-40, 0.3% Triton X-100, 3% glycerol, 1 mM DTT) supplemented with protease inhibitors (Complete Mini, Roche). The protein concentrations of whole-cell lysates were determined with the Pierce™ Rapid Gold BCA Protein Assay Kit. 30 µg of whole-cell lysates were resolved by SDS-PAGE and transferred to polyvinylidene difluoride (PVDF) membranes (Bio-Rad). Membranes were blocked with 5% non-fat dry milk in TBST (Tris-buffered saline with 0.05% TWEEN-20, pH 8.0) and probed with indicated antibodies. Immunoreactive proteins were visualized using enhanced chemiluminescence detection substrate (Amersham ECL Prime Western Blotting Detection Reagent RPN2232) using Molecular Imager ChemiDoc XRS+ System (BioRad).

For immunoprecipitation, two milligrams of lysate were incubated with the indicated antibody-conjugated beads overnight at 4 °C. The immunoprecipitants were washed twice with lysis buffer (50 mM HEPES pH 7.5, 100 mM KCl, 2 mM EDTA, 0.3% NP-40, 0.3% Triton X-100, 3% glycerol, 1 mM DTT) and thrice with tris-buffered saline (TBS). We used epitope peptide competition, acid elution and loading dye elution depending on the experiment. FLAG-tagged PRMT1 ubiquitin assays employed acid elution, PRMT1 acetylation assays used loading dye followed by boiling for elution, and epitope peptide competition of FLAG was used for elution of FLAG-tagged PRMT1 for PTM profiling using LC-MS/MS. For Myc-tagged and HA-tagged proteins, loading dye followed by boiling was used for elution. For immunoblotting, the proteins were resolved by SDS-PAGE and followed by immunoblotting analysis with indicated antibodies.

### CO-IP LC-MS/MS for interactome of PRMT1 and FBXL17

#### Immunoprecipitation of Flag-tagged PRMT1 and V5-tagged FBXL17

For the PRMT1 interactome, HEK293T cells were transfected with FLAG-PRMT1 using PolyJet (SignaGen) and cells were harvested 36 hours post-transfection in lysis buffer (50 mM HEPES pH 7.5, 100 mM KCl, 2 mM EDTA, 0.3% NP-40, 0.3% Triton X-100, 3% glycerol, 1 mM

DTT). Whole cell lysate was immunoprecipitated with anti-FLAG M2 affinity gel (Sigma-Aldrich). After overnight incubation at 4 °C, beads were washed three times with lysis buffer followed by two washes with chilled TBS. Proteins were eluted via the addition of FLAG peptide (final concentration of 150 ng/μl) and rocking for 4 hours at 4 °C. For FBXL17 interactome studies, HEK293T cells were transfected with V5-FBXL17 using PolyJet (SignaGen). Cells were harvested 36 hours post-transfection in lysis buffer. Whole cell lysate was immunoprecipitated with anti-V5 magnetic beads (CST 31628S) rocking overnight at 4 °C. After incubation, resin was washed once with lysis buffer followed by two washes with chilled TBS. For the FBXL17 interactome, on-resin digestion was performed as described below.

#### Reduction, alkylation and trypsin digestion / on-bead digestion

Equal amounts of protein were taken from both FLAG-PRMT1 and FBXL17-V5 immunoprecipitation (eluent or resin respectively) for digestion. The protein samples were reduced and alkylated using 5 mM dithiothreitol (DTT) and 10 mM iodoacetamide (IAA), respectively. Trypsin digestion was carried out using modified sequence-grade trypsin (Promega, Madison, WI) in 1:50 enzyme:protein ratio and reactions were left shaking at 37 °C overnight. Digested peptides were cleaned using C18 StageTip, vacuum dried, and stored at -20 °C before data acquisition.

#### LC-MS/MS Analysis

Thermo Scientific Orbitrap Exploris 240 mass spectrometer (ThermoFisher Scientific, Bremen, Germany) connected to the Thermo Scientific UltiMate 3000 HPLC nanoflow liquid chromatography system (ThermoFisher Scientific) was used for data acquisition. Peptide digests were reconstituted in 0.1 % formic acid in water (solvent A) to a final peptide concentration of 500 ng/μl and separated on an analytical column (75 µm × 15 cm) at a flow rate of 300 nL/min using a step gradient of 1–25 % solvent B (0.1 % formic acid in acetonitrile) for the first 100 minutes, 25–30 % for 5 minutes, 30-70 % for 5 minutes, 70–1 % for 5 minutes and 1 % for 5 minutes for a total run time of 120 minutes. The mass spectrometer was operated in data-dependent acquisition mode. A survey full scan MS (from m/z 400–1600) was acquired in the Orbitrap with a resolution of 6000 Normalized AGC target 300. Data was acquired in topN with 20 dependent scans. Peptides were fragmented using normalized collision energy with 37 % and detected at a mass resolution of 30,000. Dynamic exclusion was set for 8 s with a 10 ppm mass window.

#### Data Analysis

The raw files obtained after data acquisition were searched using Proteome Discoverer software suite version 3.0 (ThermoFisher Scientific). Data was searched against the Human Uniprot database along with known mass spectrometry contaminants using SEQUEST. Search parameters included carbamidomethylation of cysteine as static modifications. Dynamic modifications included oxidation of methionine and acetylation at protein N-terminus. The minimum peptide length was set as seven amino acids with one missed cleavage allowed. Mass tolerance was set to 10 ppm at the MS level and 0.05 Da for the MS/MS level, and the false discovery rate was set to 1% at the PSM level.

#### Detection of PRMT1 acetylation sites

HEK293T cells were transfected with FLAG-PRMT1 and Myc-p300 using PolyJet (SignaGen). Thirty-six hours post-transfection, 293T cells were treated with deacetylatase inhibitor cocktail (HY-K0030) for 1 h to block lysine deacetylation before they were harvested in lysis buffer (50 mM HEPES pH 7.5, 100 mM KCl, 2 mM EDTA, 0.3% NP-40, 0.3% Triton X-100, 3% glycerol, 1 mM DTT with protease and deacetylase inhibitors). The whole-cell lysates were immuno-precipitated with ANTI-FLAG® M2 affinity gel (Sigma-Aldrich), in the presence of deacetylase inhibitor cocktail. Beads were washed twice with lysis buffer followed by three washes with chilled TBS. FLAG-tagged PRMT1 was eluted from beads using FLAG peptide competition (Sigma-Aldrich). Peptide was added at a 150 ng/µl concentration and allowed to incubate for four hours at 4°C. Protein samples were reduced with dithiothreitol, and cysteine residues were alkylated with iodoacetamide as described above. Samples were then digested overnight at 37 °C using Trypsin/LysC enzyme. Peptide mixtures were cleaned using a C18 ziptip column and injected on a Thermo Scientific UltiMate 3000 HPLC nanoflow liquid chromatography system coupled to a Thermo Scientific Orbitrap Exploris 240 mass spectrometer using a 75mm i.d 15 cm C18 microcapillary column at a flow rate of 300 nL/ min. Data dependent MS/MS acquisitions were performed using the Top 13 method. All MS/MS samples were analyzed using SEQUEST. SEQUEST was set up to search the Human PRMT1 sequence assuming the digestion enzyme trypsin, and acetylation of lysine was specified as a variable modification.

#### Detection of PRMT1 ubiquitination sites

HEK293T cells were transfected with FLAG-PRMT1 using PolyJet (SignaGen). Thirty-six hours post-transfection, cells were treated with MG-132 for 6 hours. Thirty minutes before harvesting, cells were treated with deubiquitinase (DUB) inhibitor PR-169 and lysed in denaturing buffer (50 mM HEPES pH 7.4, 100 mM KCl, 2mM EDTA, 0.3% NP-40, 0.3% Triton X-100) containing protease and deubiquitinase inhibitors. The whole-cell lysates were immunoprecipitated with ANTI-FLAG® M2 affinity gel (Sigma-Aldrich) in the presence of PR-169. Protein elution was done using FLAG-peptides (Sigma-Aldrich). The protein sample was then reduced with DTT, and cysteine residues were alkylated with iodoacetamide, as described above. The protein was digested overnight at 37 °C using the Trypsin/LysC enzyme. Peptide mixtures were cleaned using a C18 ziptip column and injected on a Thermo Scientific UltiMate 3000 HPLC nanoflow liquid chromatography system coupled to a Thermo Scientific Orbitrap Exploris 240 mass spectrometer using a 75mm i.d 15 cm C18 microcapillary column at a flow rate of 300 nL/ min. Data dependent MS/MS acquisitions were performed using the Top 13 method. All MS/MS samples were analyzed using SEQUEST. SEQUEST was set up to search the Human PRMT1 sequence assuming the digestion enzyme trypsin. Iso-peptides of GG and LRGG at lysine were specified as variable modifications.

#### Cycloheximide chase assays

For endogenous PRMT1, HEK293T cells were seeded at a density of 5 × 10^5^ cells per well in a 6 well plate. At 80% confluency, cells were treated with 50 μg/ml of cycloheximide. Cells were harvested at 5, 10, and 15 hours after CHX treatment. For stably expressed FLAG-PRMT1, the relevant Flp-In HEK293T cells were seeded at a density of 5 × 10^5^ cells per well in a 6 well plate. Four hours after seeding, cells were treated with tetracycline to induce PRMT1 expression. Once cells reached 80% of confluency, treatment with 50 μg/ml of CHX was started and cells were harvested at 5, 10, and 15 hours after treatment. For overexpression, HEK293T cells were seeded at a density of 5 × 10^5^ cells per well in 6 well plate and transfected with FLAG-PRMT1 using PolyJet (SignaGen) at 90% confluency. Cells were trypsinized and plated at low density (5 × 10^4^) 48 hours post-transfection and were allowed to grow for 72 hours. After 72 hours, 50 μg/ml of cycloheximide was added and cells were harvested at 5, 10, and 15 hours after treatment.

#### In-cell ubiquitination assays

HEK293T cells were co-transfected with FLAG-PRMT1 constructs along with HA-tagged ubiquitin (HA-Ub) using PolyJet (SignaGen). At 48 hours post-transfection, cells were treated with MG-132 for 3 hours and deubiquitinase inhibitor (PR-619) for 15 minutes to inhibit DUBs before harvesting. Cells were lysed via sonication in lysis buffer (50 mM HEPES pH 7.5, 100 mM KCl, 2 mM EDTA, 0.3% NP-40, 0.3% Triton X-100, 3% glycerol, 1 mM DTT) supplemented with protease inhibitor cocktail (A32955) and DUB inhibitor (PR-619). After overnight incubation with ANTI-FLAG M2 Affinity Gel (Sigma-Aldrich), beads were washed three times with lysis buffer followed by three washes with chilled PBS. Proteins were eluted under acidic conditions. Eluents were resolved by SDS-PAGE and immunoblotted using indicated antibodies.

#### In-cell acetylation assays

Detection of acetylation within cells were performed as follows. HEK293T cells were grown in 10-cm dish. Once cells reached 90% confluency, cells were co-transfected with Myc-p300 (acetyltransferase) and FLAG-PRMT1. Media was changed to standard media 12 hours post-transfection and cells were allowed to grow for 48 hours. Before harvesting, cells were treated with a deacetylase inhibitor cocktail for 30 minutes. Cells were lysed via sonication in lysis buffer (50 mM HEPES pH 7.5, 100 mM KCl, 2 mM EDTA, 0.3% NP-40, 0.3% Triton X-100, 3% glycerol, 1 mM DTT) buffer supplemented with protease inhibitor cocktail (A32955) and deacetylase inhibitor cocktail (MEC HY-K0030). PRMT1 was immunoprecipitated using ANTI-FLAG M2 Affinity Gel (Sigma-Aldrich) and eluted using FLAG peptides (Sigma-Aldrich). Samples were immunoblotted for anti-acetyl lysine to look for the overall acetylation status of PRMT1.

#### Cloning

In general, plasmids were maintained in DH5α cells (NEB), with exceptions noted. For pcDNA3.1-PRMT1 and variants, a gene fragment for human PRMT1 with a C-terminal FLAG epitope tag was inserted into a pcDNA3.1 vector (Genscript). From this construct, all point mutants of PRMT1 at K228 and K233 were made by site-directed mutagenesis (Q5 Site Directed Mutagenesis kit, NEB). For downstream stable cell line generation, PRMT1-FLAG was subcloned from pcDNA3.1 by PCR amplification and inserted into a pcDNA5 backbone (Thermo Scientific) using Gibson Assembly (NEB). FBXL17 constructs were created using a gene fragment corresponding to human FBXL17 (DNASU) and inserting this into a pcDNA3.1 vector using Gibson Assembly (NEB). For downstream stable cell line generation, FBXL17-V5 was amplified by PCR and inserted into a pcDNA5 backbone (Thermo Scientific) using Gibson Assembly (NEB). pRK5-HA-Ubiquitin constructs were a gift from Craig Crews, Yale University.

For the genetic code expansion constructs, the initial E312 and E336 plasmids were a gift from Dr. Jason Chin. To generate plasmids for our study, human PRMT1 was amplified and inserted into a linearized E336 plasmid using Gibson Assembly (NEB). The necessary TAG mutants, K228TAG and K233TAG, were subsequently made by site-directed mutagenesis (Q5, NEB). The completed E336-PRMT1 variants include PRMT1 under EF-1α control, as well as 4 copies of the Mb tRNA under U6 promoter. The E336-PRMT1 variants were maintained in Stbl3 cells (NEB) and grown at 30 .

##### Relevant amino acid sequences

Human PRMT-FLAG (FLAG in bold, K228 and K233 residues in bold): MAAAEAANCIMENFVATLANGMSLQPPLEEVSCGQAESSEKPNAEDMTSKDYYFDSYAHFGIHE EMLKDEVRTLTYRNSMFHNRHLFKDKVVLDVGSGTGILCMFAAKAGARKVIGIECSSISDYAVK IVKANKLDHVVTIIKGKVEEVELPVEKVDIIISEWMGYCLFYESMLNTVLYARDKWLAPDGLIF PDRATLYVTAIEDRQYKDYKIHWWENVYGFDMSCI**K**DVAI**K**EPLVDVVDPKQLVTNACLIKEVD IYTVKVEDLTFTSPFCLQVKRNDYVHALVAYFNIEFTRCHKRTGFSTSPESPYTHWKQTVFYME DYLTVKTGEEIFGTIGMRPNAKNNRDLDFTIDLDFKGQLCELSCSTDYRMR**DYKDDDDK**

##### Human Fbxl17 - V5 (V5 in bold)

MGHLLSKEPRNRPSQKRPRCCSWCRRRRPLLRLPRRTPAKVPPQPAAPRSRDCFFRGPCMLCFI VHSPGAPAPAGPEEEPPLSPPPRDGAYAAASSSQHLARRYAALAAEDCAAAARRFLLSSAAAAA AAAASASSPASCCKELGLAAAAAWEQQGRSLFLASLGPVRFLGPPAAVQLFRGPTPSPAELPTP PEMVCKRKGAGVPACTPCKQPRCGGGGCGGGGGGGGGGGPAGGGASPPRPPDAGCCQAPEQPPQ PLCPPPSSPTSEGAPTEAGGDAVRAGGTAPLSAQQQHECGDADCRESPENPCDCHREPPPETPD

INQLPPSILLKIFSNLSLDERCLSASLVCKYWRDLCLDFQFWKQLDLSSRQQVTDELLEKIASR SQNIIEINISDCRSMSDNGVCVLAFKCPGLLRYTAYRCKQLSDTSIIAVASHCPLLQKVHVGNQ DKLTDEGLKQLGSKCRELKDIHFGQCYKISDEGMIVIAKGCLKLQRIYMQENKLVTDQSVKAFA EHCPELQYVGFMGCSVTSKGVIHLTKLRNLSSLDLRHITELDNETVMEIVKRCKNLSSLNLCLN WIINDRCVEVIAKEGQNLKELYLVSCKITDYALIAIGRYSMTIETVDVGWCKEITDQGATLIAQ SSKSLRYLGLMRCDKVNEVTVEQLVQQYPHITFSTVLQDCKRTLERAYQMGWTPNMSAASS**MGK PIPNPLLGLDST**

Human HA - Ubiquitin (HA in bold): MG**YPYDVPDYA**DLNGGGGGSTMQIFVKTLTGKTITLEVEPSDTIENVKAKIQDKEGIPPDQQRL IFAGKQLEDGRTLSDYNIQKESTLHLVLRLRGG

#### Genetic code expansion

For genetic code expansion experiments, we transfected the stable HEK293T + E312 cells with cloned E336-PRMT1 variants, containing 4 more copies of the tRNA and the PRMT1 variant of interest, using PolyJet (SignaGen) and immediately added 10 mM Nε-acetyl-lysine (Sigma-Aldrich) to the media. Upon harvesting, deacetylase inhibitor cocktail (CST 362323) was added to lysis buffer at a 100-fold dilution.

## Results

Seeking to identify protein-protein interactions that may play a role in controlling PRMT1 regulation, we initially performed a co-immunoprecipitation mass spectrometry analysis in HEK293T cells (Fig. 1A). We identified 693 high confidence interactors, defined as interactors found across three replicates, which comprised 450 known interactors and 243 new interaction partners (Fig. 1B, Table S1). High confidence interactors known to be “PTM writer” enzymes, or enzymes known to attach PTMs onto target proteins, were of particular interest due to their known regulatory function (Table S1). Intriguingly, acetyltransferase p300 was identified as a high confidence interaction partner of PRMT1 and vice versa under p300 co-IP LC-MS/MS (Table S2). Acetylation of PRMT1 has only been reported once (Lai et al., 2017) and is therefore under-characterized as a PRMT1 regulatory mechanism. The p300-PRMT interaction was verified through co-immunoprecipitation experiments in HEK293T cells overexpressing Myc-tagged p300 (Myc-p300) (Fig. 1C). As an additional control, we performed the reverse experiment and demonstrated that FLAG-tagged PRMT1 (FLAG-PRMT1) was equally able to co-immunoprecipitate endogenous p300 (Fig. 1C). Furthermore, we demonstrated that the p300-PRMT1 interaction resulted in acetylation of PRMT1 through the increase of acetyl-lysine signal on an immunoprecipitated FLAG-PRMT1 following co-expression of Myc-p300 and FLAG-PRMT1. Further, this acetylation can be abrogated by inhibition of p300 acetyltransferase activity using A-485 (Lasko et al., 2017) (Fig. 1D). Having demonstrated p300-catalyzed acetylation of PRMT1, we performed a detailed characterization of the acetylation and ubiquitination sites of PRMT1 (Fig. 1E) in order to build a more complete picture on the impact of lysine acetylation on PRMT1 homeostasis. We identified many known acetylation (Fig. S1B/C, Table S3) and ubiquitination (Table S4) sites but also revealed additional novel sites, including several with both acetylation and ubiquitination in equilibrium, suggesting the interplay of acetylation and ubiquitination may play a role in regulating PRMT1 stability.

**Figure 1.**
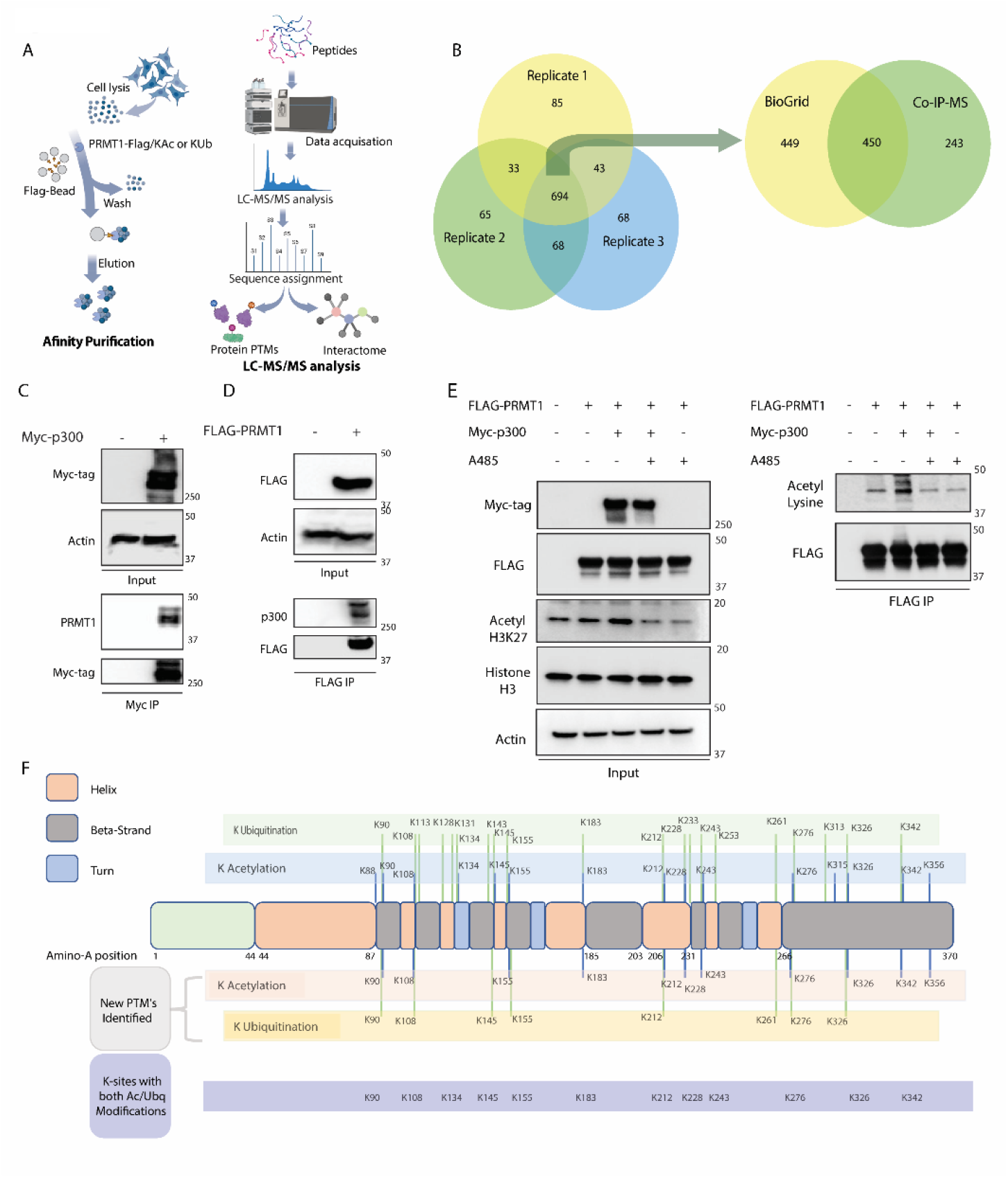
PRMT1 Interactome and Lysine Acetylation reveal p300 mediated acetylation interfaces with ubiquitination. (A) Experimental workflow (B) Venn diagram showing interactome of PRMT1 from Co-IP-LC-MS/MS compared with BioGrid data. (C) Western blotting validating the interaction of p300 with PRMT1, Myc-tagged p300 was overexpressed and immunoprecipitated. (D) Western blotting validating interaction of PRMT1 with p300, FLAG-tagged PRMT1 was overexpressed and immunoprecipitated using anti-FLAG affinity gel. (E) IP-western blotting showing acetylation of PRMT1 in p300 overexpressed and inhibited conditions. (F) Acetylation and ubiquitination sites of PRMT1 identified using IP-LC-MS/MS.

During the course of these experiments, we noticed an intriguing phenomenon; we consistently observed reduced equilibrium expression of PRMT1 when p300 was overexpressed and conversely not when p300 was inhibited (Fig. 2A). Using an existing RNA-Seq data set, we investigated if this observation could be explained by altered transcriptional regulation but saw no effect of p300 overexpression or inhibition on PRMT1 expression at the transcript level (Fig. 2B). The same phenomenon was observed with overexpressed FLAG-PRMT1, ruling out any effect of PRMT1-modulated chromatin dynamics (Fig. 2C). This led us to hypothesize that p300-mediated acetylation could be affecting PRMT1 stability post-translationally, further hinting at the interplay of acetylation and ubiquitination. To test this hypothesis, we performed cycloheximide chase experiments in HEK293T cells to observe PRMT1 half-life under p300 overexpression/inhibited conditions (Fig. 2D). Similar to the literature (Luo et al., 2024), we observed that PRMT1 has a relatively long half-life of approximately 15 hours when unperturbed, which is significantly shortened by p300 overexpression and extended by p300 inhibition (Fig. 2D). This phenotype was conserved between endogenous PRMT1 (Fig. 2D) and ectopically expressed FLAG-PRMT1 (Fig. 2E), again demonstrating that results were not reliant on chromatin dynamics. We next looked to ensure that changes in PRMT1 stability were enacted through the ubiquitin-proteasome system through assessing the ability of FLAG-PRMT1 to pull down a co-transfected HA-ubiquitin (HA-Ub) construct under proteasome-inhibited conditions. We demonstrated that PRMT1 is robustly ubiquitinated (Fig. 2F) and that this ubiquitination is significantly increased by p300 overexpression (Fig. 2G). Further, we showed that enhanced ubiquitination is dependent on the acetyltransferase activity of p300 as inhibition of p300s acetyltransferase domain with inhibitor A-485 restored the control levels of ubiquitination (Fig. 2G).

**Figure 2.**
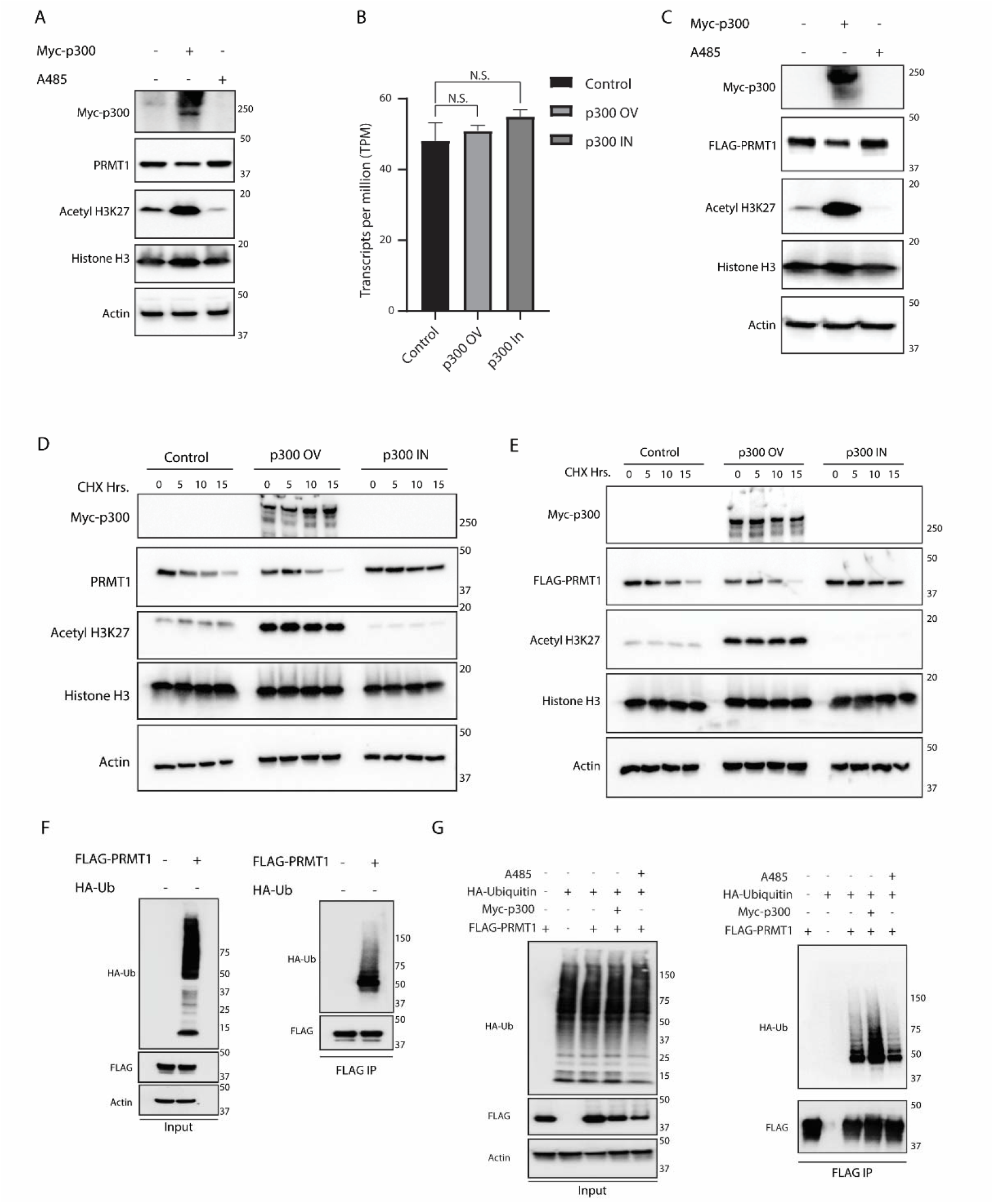
p300-mediated acetylation of PRMT1 decreases PRMT1 half-life and increases PRMT1 ubiquitination. (A) Western blotting showing expression of endogenous PRMT1 across p300 overexpression/inhibited conditions. (B) mRNA expression of PRMT1 in p300 overexpressed and p300 inhibited conditions. (N.S. – not significant) (C) Western blotting showing expression of overexpressed FLAG-PRMT1 across p300 overexpression/inhibited conditions. (D) Western blotting showing half-life of endogenous PRMT1 across p300 overexpression/inhibited conditions under cycloheximide treatment. Cells were treated with 50 μg/ml of cycloheximide. (E) Western blotting showing half-life of overexpressed PRMT1 across p300 overexpression/inhibited conditions under cycloheximide treatment. Cells were treated with 50 μg/ml of cycloheximide. (F) IP-Western blotting showing the ubiquitination of PRMT1, Flag-tagged PRMT1 was co-expressed with HA-tagged ubiquitin followed by FLAG-IP. (G) Western blotting illustrates ubiquitination of FLAG-PRMT1 across p300 overexpressed/inhibited conditions. FLAG-tagged PRMT1 was co-expressed with HA-tagged ubiquitin, FLAG-PRMT1 was immunoprecipitated and samples were probed with anti-HA antibody.

Previous literature (Lai *et al*., 2017) suggested that the degradation of PRMT1 may be mediated by the SCF^FBXL17^ complex. SCF (SKP1-cullin 1-F box) complexes are comprised of a central cullin-1 scaffold that binds RBX1 and SKP1 on its N- and C-terminal ends. RBX1 and SKP1 function as adaptor proteins that bind E2s and F-box protein:target protein complexes, respectively, to bring them into proximity for ubiquitin transfer. F-box proteins are the only subunit of the SCF complex that directly bind the target protein and are therefore the major determinants of ubiquitination targets. PRMT1s ability to interact with FBXL17 (F-box protein leucine rich repeat 17) could therefore have direct implications on PRMT1 homeostasis. Indeed, an FBXL17 co-IP LC-MS/MS (Fig. S2A, Table S6) suggested PRMT1 as a high confidence interactor. We proceeded to confirm the interaction between PRMT1 and FBXL17 through co-immunoprecipitation with a V5-tagged FBXL17 construct (V5-FBXL17) (Fig. 3A). Both endogenous PRMT and Flag-PRMT1 were co-immunoprecipitated by V5-FBXL17. We also validated the functionality of this interaction by overexpressing V5-tagged FBXL17 and observing a reduction in levels of endogenous PRMT1 in HEK293T cells (Fig. S2B) and exogenous PRMT in HEK293T FlpIn^Flag-PRMT1^ Cells (Fig. 3B/C). Additionally, we generated a FBXL17 KO cell line using CRISPR/Cas9 (Fig. 3D) to further validate the role of FBXL17 in PRMT1 turnover. Cycloheximide chase experiments using V5-FBXL17 overexpression and the FBXL17 KO cells show that FBXL17 knock out significantly extends the half-life of endogenous PRMT1 and V5-FBXL17 overexpression decreases PRMT1 half-life even when compared to unperturbed conditions (Fig. 3E). Finally, to confirm that FBXL17-mediated changes in PRMT1 half-life are enacted through the ubiquitin-proteasome system, we again used co-transfected HA-Ub to demonstrate that PRMT1 ubiquitination is depleted in FBXL17 KO cells (Fig. 3F). We also confirmed that PRMT1 ubiquitination is dominated by K48-linked ubiquitin by demonstrating that overexpression of a K48R Ub dominant negative mutant rescues PRMT1 stability, while K63R does not (Fig. S2C-E), suggesting that the canonical K48-linked polyubiquitin signal is responsible for PRMT1 degradation.

**Figure 3.**
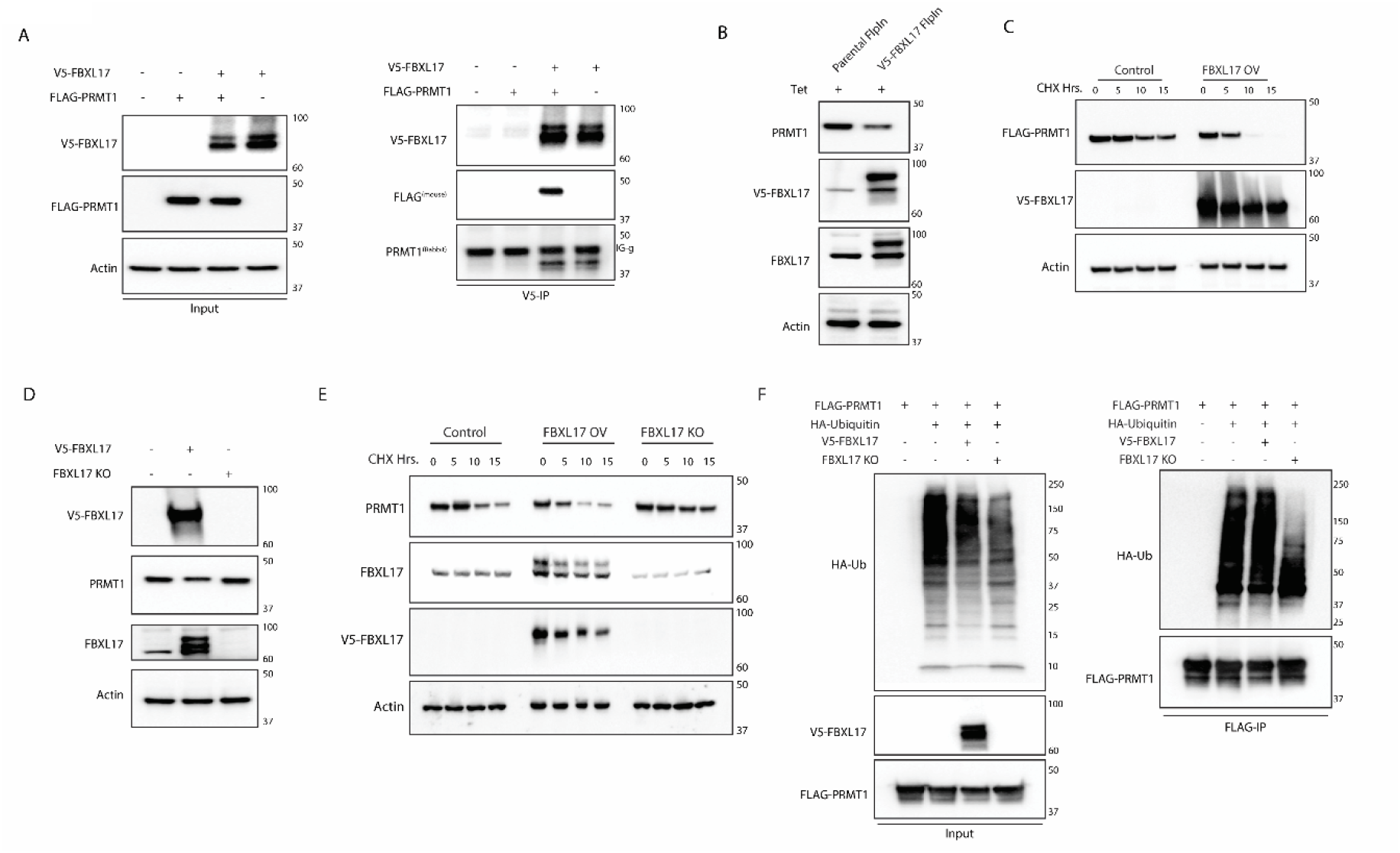
FBXL17 interaction with PRMT1 affects PRMT1 half-life and ubiquitination. (A) Co-immunoprecipitation and immunoblotting validating the interaction of FBXL17 and PRMT1. V5-FBXL17 was overexpressed both alone and with FLAG-tagged PRMT1, followed by immunoprecipitation of V5-tagged FBXL17 (B) Western blotting shows expression of FLAG-PRMT1 in control and FBXL17 overexpressed conditions. (C) Western blotting showing half-life of FLAG-PRMT1 under cycloheximide chase conditions (50 µg/ml) in control and FBXL17 overexpressed conditions. (D) Validation of FBXL17 KO cell lines. (E) Western blotting showing the half-life of endogenous PRMT1 in control, FBXL17 overexpressed and FBXL17 KO conditions (F) Western blotting showing ubiquitination of PRMT1 in control, FBXL17 overexpressed and KO conditions. FLAG-PRMT1 was co-expressed with HA-tagged ubiquitin followed by FLAG-tag immunoprecipitation and HA-tag immunoblotting.

Having demonstrated that both p300-mediated acetylation and FBXL17 affect PRMT1 half-life and ubiquitination, we next sought to identify how PRMT1 acetylation and the FBXL17 interaction may be linked. We generated Lysine (K) -to-Asparagine (Q) mutants as acetylation mimics and K-to-Arginine (R) mutants as non-acetylatable lysine mimics at PRMT1 acetylation sites identified by in our PTM analysis (Fig. 1E) and previous literature (Lai et al., 2017), before characterizing them using cycloheximide chase experiments. Mutations at K228 and K233 had the highest impact on PRMT1 half-life (Fig. 4A). The effect of the K228Q mutant was particularly striking as it drastically reduced PRMT1 half-life, while the K228R mutant extended it. Concurrent or independent mutation of the adjacent K233 residue had little effect, suggesting that this is a specific interaction that can be altered by this residue-specific acetylation rather than global protein stability (Fig. 4A). We also demonstrated that PRMT1^K228Q^, possessing the shortest half-life, is heavily ubiquitinated whilst PRMT1^K228R/K233Q^, possessing the longest half-life, has significantly reduced ubiquitination (Fig. 4B). Furthermore, we demonstrate by co-immunoprecipitation between FLAG-PRMT1 and V5-FBXL17 that the enhanced turnover and ubiquitination of PRMT1 correlates with the ability of PRMT1 to interact with FBXL17 (Fig. 4C), as PRMT1^K228Q^ containing variants demonstrated the strongest interaction with FBXL17.

**Figure 4.**
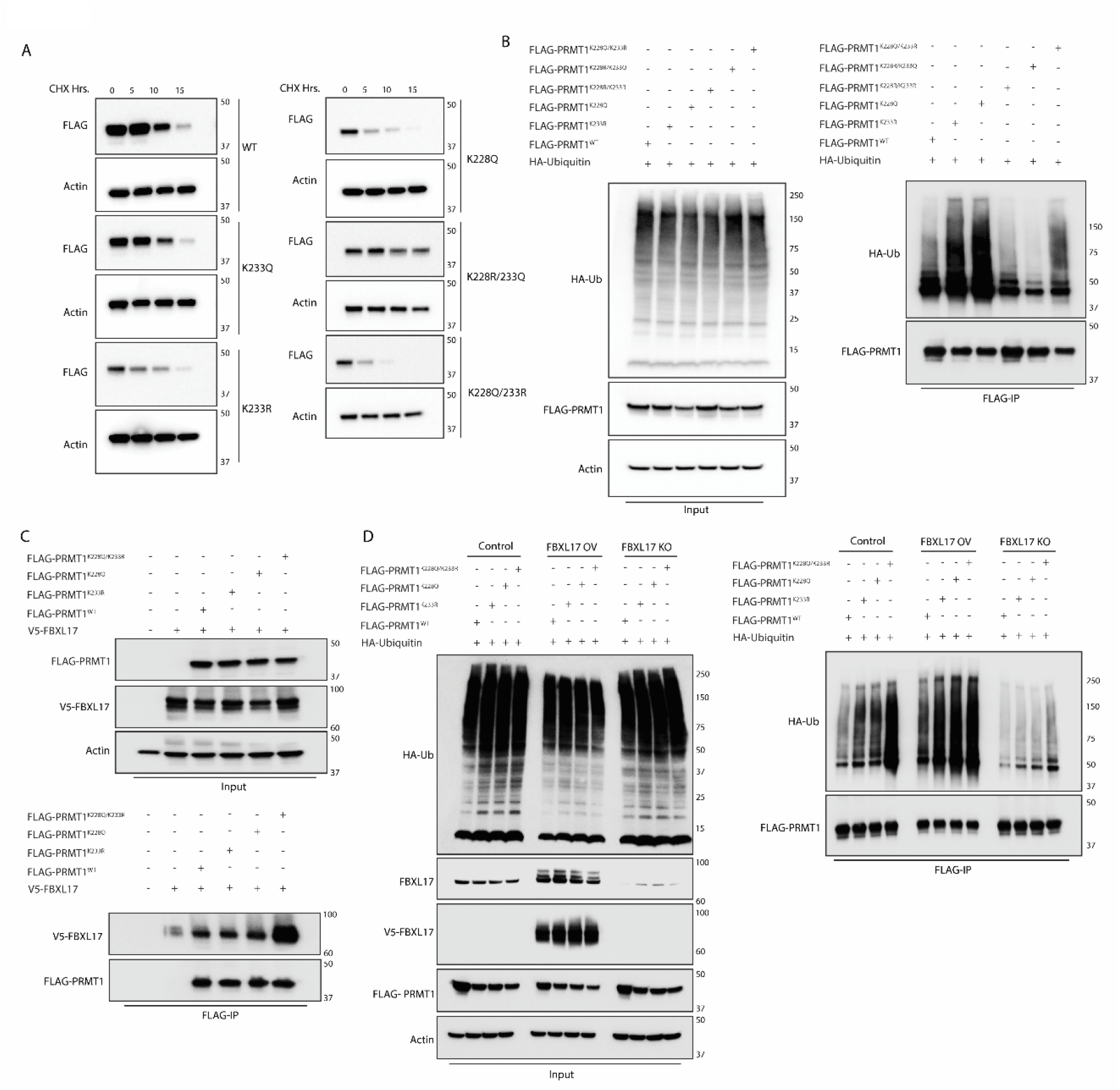
PRMT1 acetylation at K228 increases FBXL17 interaction and ubiquitin-mediated degradation. (A) Western blotting showing the half-life of PRMT1 mutants under cycloheximide treatment (50 μg/ml). FLAG-tagged PRMT1 mutants were overexpressed. (B) Western blotting showing the ubiquitination of PRMT1 variants, FLAG-tagged PRMT1 was co-expressed with HA-tagged ubiquitin followed by anti-FLAG-immunoprecipitation. (C) Western blotting showing the interaction of FBXL17 with PRMT1 mutants. FLAG-PRMT1 mutants were co-expressed with V5-tagged FBXL17 followed by FLAG co-immunoprecipitation. (D) Western blotting showing ubiquitination of PRMT1 mutants in FBXL17 overexpressed and knockout conditions. FLAG-PRMT1 mutants were co-expressed with HA-tagged ubiquitin in the indicated cell lines, followed by FLAG-immunoprecipitation.

Importantly, the ubiquitination of these PRMT1 variants was enhanced by FBXL17 overexpression and reduced in our FBXL17 KO cells (Fig. 4D), as were the half-lives under cycloheximide chase conditions (Figure S3), thus confirming that installing acetylation mimics at specific sites can drastically change PRMT1’s ability to bind FBXL17, ubiquitination profile, and rate of degradation.

However, acetyl-lysine and non-acetylatable lysine mimicry via Q/R mutations do not represent true acetylation marks and may accordingly confound results. To address this problem and unequivocally test the hypothesis that acetylation of PRMT1 at K228 results in its FBXL17-mediated degradation, we employed a mammalian genetic code expansion (GCE) approach originally reported by Chin and co-workers (Elsässer et al., 2016) to site-specifically incorporate authentic acetylated lysine (AcK) residues at K228 or K233 (Fig. 5A). HEK293T cells were engineered via PiggyBac transposase (Li et al., 2013) to express a orthogonal amino-acyl tRNA synthetase/tRNA system evolved to incorporate AcK residues at an amber stop codon (TAG) sites. Engineered cells were then transfected with PRMT1 plasmids encoding a TAG codon at the relevant sites. The inclusion of a FLAG epitope tag at the *C*-terminus ensures that a FLAG immunoreactive band is present only when the non-canonical amino acid is successfully incorporated, resulting in full-length PRMT1. The addition of the GCE machinery has no effect on the expression of PRMT1 from a plasmid encoding PRMT1^WT^ but enables the expression of PRMT1^228TAG^ or PRMT1^233TAG^ through incorporating the non-canonical AcK residue at the TAG codon site (Fig. 5B). Small amounts of truncation product are visible on the PRMT1 blot since the PRMT1 antibody recognizes an epitope upstream of the TAG sites but are not visible in the c-terminal FLAG immunoblots (Fig. 5B). This approach was verified by our demonstration that full-length FLAG-tagged protein is dependent upon the addition of exogenous AcK to the media (Fig. 5C). Furthermore, FLAG immunoprecipitation of cells transfected with FLAG-PRMT1^228TAG^ or FLAG-PRMT1^233TAG^ displays robust anti-AcK immunoreactivity (Fig. 5C). Crucially, the incorporation of the AcK is site specific, as demonstrated by IP-LC-MS/MS analysis of tryptic peptides followed by label free quantification of the detected peptides across samples (Fig. 5D/E). K228Ac peptides are predominantly detected in PRMT1^228TAG^ transfected cells as anticipated and not in PRMT1^233TAG^, PRMT1^WT^ or mock transfected cells (Fig. 5D). K233Ac peptides are likewise only detected in PRMT1^233TAG^ cells (Fig. 5E). Following the validation of this system, we probed the effect of site-specific lysine acetylation on PRMT1 stability using cycloheximide chase experiments (Fig. 5F). Consistent with our hypothesis, the FLAG signal is depleted much more rapidly in the cells expressing PRMT1^228TAG^ than either PRMT1^WT^ or PRMT1^233TAG^ (Fig. 5F). These findings demonstrate that acetylation of K228 is responsible for the degradation of PRMT1 under these otherwise non-perturbing conditions. Finally, we demonstrated that PRMT1^228TAG^ has a significantly increased ability to bind FBXL17, thus explaining the reduced half-life (Fig. 5G).

**Figure 5.**
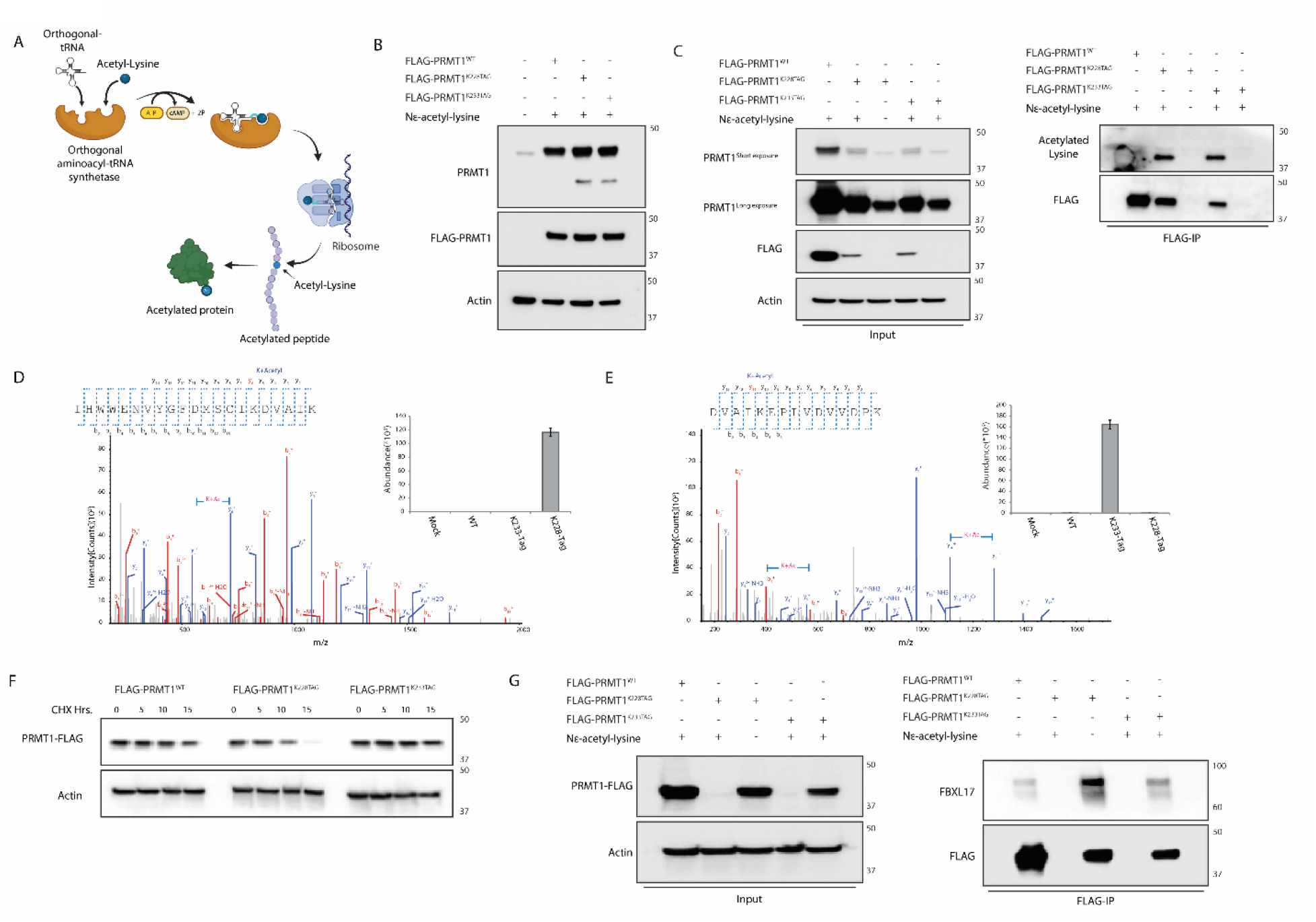
Authentic acetyl-lysine incorporation via genetic code expansion shows decreased half-life of K228 acetylated PRMT1. (A) Schematic of genetic code expansion approach. (B) Western blotting validating PRMT1 genetic code expansion. (C) Immunoprecipitation reveals conditional incorporation of acetylated lysine. FLAG-PRMT1^WT^, PRMT1^228TAG^ or PRMT1^233TAG^ were transfected followed by N-acetyl-lysine treatment. FLAG-immunoprecipitation followed by immunoblotting confirms incorporation of acetylated lysine. (D) MS/MS spectra of peptide with acetylated K228 and bar graph shows the abundance of acetylated peptide across the conditions. (E) MS/MS spectra of peptide with acetylated K233 and bar graph shows the abundance of acetylated peptide across the conditions. (F) Western blotting showing the half-life of PRMT1 (WT, 228TAG and 233TAG) GCE variants under cycloheximide treatment (50 μg/ml). (G) Co-immunoprecipitation of acetylated PRMT1 GCE variants reveals PRMT1^228TAG^ has an enhanced interaction with FBXL17.

## Discussion

Intrigued by the initial observation that PRMT1 protein levels may be modulated by acetylation flux in cells, we sought to understand the post-translational regulation of this highly disease relevant protein. Using a series of mass-spectrometry based proteomics and cellular biochemistry experiments, we determined that p300 acetylates PRMT1 which triggers a cascade of events resulting in its degradation (Fig. 6). We confirmed p300 directly acetylates PRMT1 whilst having no significant impact on PRMT1 transcript levels, leading to the hypothesis that p300-mediated acetylation post-translationally regulates PRMT1 stability. Careful experiments using cycloheximide chase experiments confirmed this hypothesis and demonstrated that p300-mediated acetylation correlates with PRMT1 ubiquitination and degradation. Furthermore, we identified substrate adaptor FBXL17 and implicated it in acetyl-specific degradation of PRMT1. Our results confirm that, at both exogenous and endogenous proteins levels, FBXL17 is involved with the turnover of PRMT1. Finally, we identify K228 as the site of acetylation which is largely responsible for PRMT1’s ability to interact with FBXL17 and therefore PRMT1’s ubiquitin mediated. Mutation at the K228 site drastically alters protein half-life, concurrent with the enhanced FBXL17 interaction and subsequent ubiquitination. Our results thus provide key mechanistic insight into the post-translational control of PRMT1 stability.

**Figure 6.**
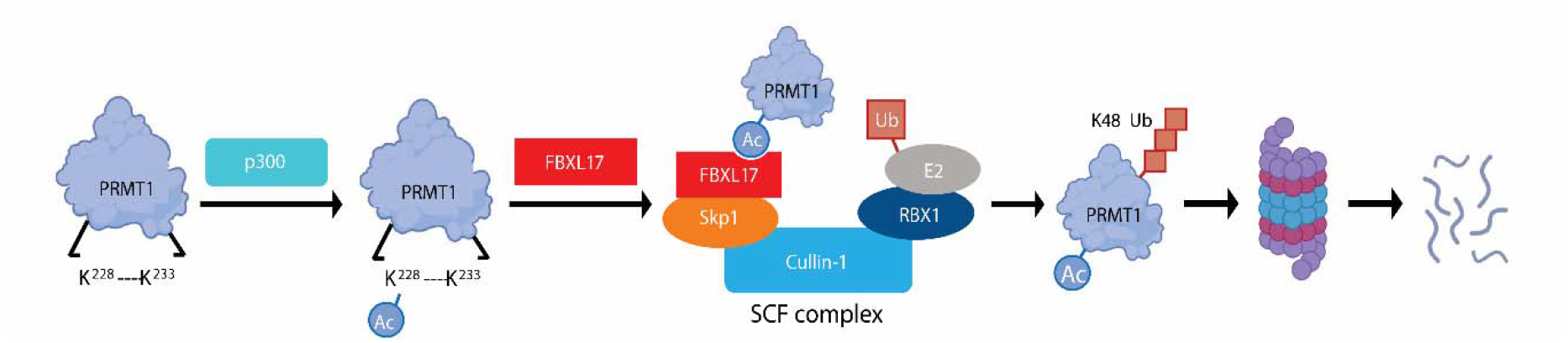
Schematic of PRMT1 Ubiquitination by FBXL17 under Acetylation Control.

PRMT1’s crucial role in transcription, signal transduction, and metabolism has been demonstrated to be a viable therapeutic target in several cancers. However, due to toxicity and lack of efficacy demonstrated by inhibitors aimed at PRMT1 catalytic activity (El-Khoueiry et al., 2023), PRMT1 therapeutic design must pivot to target other regulatory facets of PRMT1 structure and function. PTMs can play significant regulatory roles in altering target protein function, localization, interactors and/or homeostasis. The role of PTMs in PRMT1 maintenance and regulation are largely under-characterized but there is increasing evidence of phosphorylation also playing a role in PRMT1 regulation (Rust et al., 2014). Phosphorylation of Tyr291 has been shown to dramatically alter the PRMT1 interactome and generally decrease PRMT1 methyltransferase activity towards well-characterized substrates such as histone H4 and estrogen receptor alpha. Further, kinases CSNK1a1 and ULK3 have been linked to PRMT1 regulation although the exact phosphorylated sites and mechanisms are not understood (Bao et al., 2017; Goruppi et al., 2023). Outside of phosphorylation, oxidation of Cys101 and Cys208 has been shown to negatively correlate with PRMT1 methyltransferase activity (Morales et al., 2015). Finally, as noted, acetylation-mediated shifts in PRMT1 homeostasis have been previously identified (Lai et al., 2017). Our results partially conflict with these findings as we demonstrate that acetylation at K228, rather than K233, is responsible for constituting the PRMT1-FBXL17 interaction.

Given the importance of PRMT1 homeostasis to cellular arginine methylation, there are multiple mechanisms by which PRMT1 can be degraded. For example, it has also been reported to be modulated by other E3 ligases, including TRIM48 (Hirata et al., 2017) and FBXO7 (Luo *et al*., 2024). Excitingly, both the TRIM48-PRMT1 and FBXO7-PRMT1 interactions are preliminarily evidenced to be viable therapeutic targets for inhibiting oncogenic metabolism exhibited by hepatocellular carcinoma (Li *et al*., 2023; Luo *et al*., 2024). Our findings supplement the growing landscape of PRMT1 homeostasis by providing evidence for a relationship between PRMT1s p300-mediated acetylation and its FBXL17-mediated ubiquitination and degradation. We contribute a novel mechanism extensively proven by our use of both acetyl-lysine mimics and biologically authentic acetyl-lysines incorporated via genetic code expansion. Our mechanistic model (Fig. 6) therefore further elucidates the role of acetylation in PRMT1 regulation and contributes a regulatory mechanism by which PRMT1 can potentially be targeted in disease contexts.

In summary, we demonstrate an acetylation regulated degron of a key protein determinant of disease, PRMT1, and identify its cognate E3 ligase substrate adaptor, FBXL17. This provides a prototypical example of the emerging concept of acetylation regulated protein degradation and enhances the mechanistic understanding of the roles of lysine acetylation beyond chromatin. Given that lysine acetylation is the third most observed PTM in mammalian biology, it is perhaps not surprising that it plays significant roles in important processes such as protein homeostasis, but these roles remain understudied, largely due to a lack of suitable technologies. Here we utilized a genetic code expansion approach to install authentic acetylation marks site-specifically, therefore enabling the first direct evidence that site specific acetylation can induce protein degradation. This technological step forward enables the robust validation and mechanistic understanding of the impact of acetylation on protein half-life and opens new realms of exploration into the control of protein homeostasis. Notable examples, such as ACLY (Lin et al., 2013), exist where acetylation and ubiquitination are mutually exclusive at the same lysine (Choi et al., 2017; Perez-Oquendo et al., 2023; Xu et al., 2022; Yang et al., 2023; Zhang et al., 2023a). However, there are only scarce reports of lysine acetylation inducing ubiquitination (Jiang et al., 2011; Lai *et al*., 2017; Van Nguyen et al., 2016). We hypothesize that many other proteins may possess acetyl-lysine dependent degrons and undergo complex interplay with ubiquitination. Future work in our lab will focus on the mechanistic understanding of acetyl-degrons across proteins and their impact on disease.

## Acknowledgements

This work was supported by a research grant from the Leukemia Research Foundation and the National Institutes of Health R35-GM142505 (to G.M.B.). J.N.B. was supported by an NSF Graduate Research Fellowship. We are grateful to members of the Burslem lab for useful discussion. The Piggybac plasmids encoding the AcK orthogonal tRNA and amino-acyl tRNA synthetase were a kind gift from Jason Chin and the Piggybac Transposase plasmid was a kind gift from Stephanie Grainger.

## Conflicts of interest

There are no conflicts to declare.

## Notes

### Competing Interest Statement

The authors have declared no competing interest.

