## Supplemental Figures for "The Interplay of Acetylation and Ubiquitination Controls PRMT1 Homeostasis"

### Supporting Information

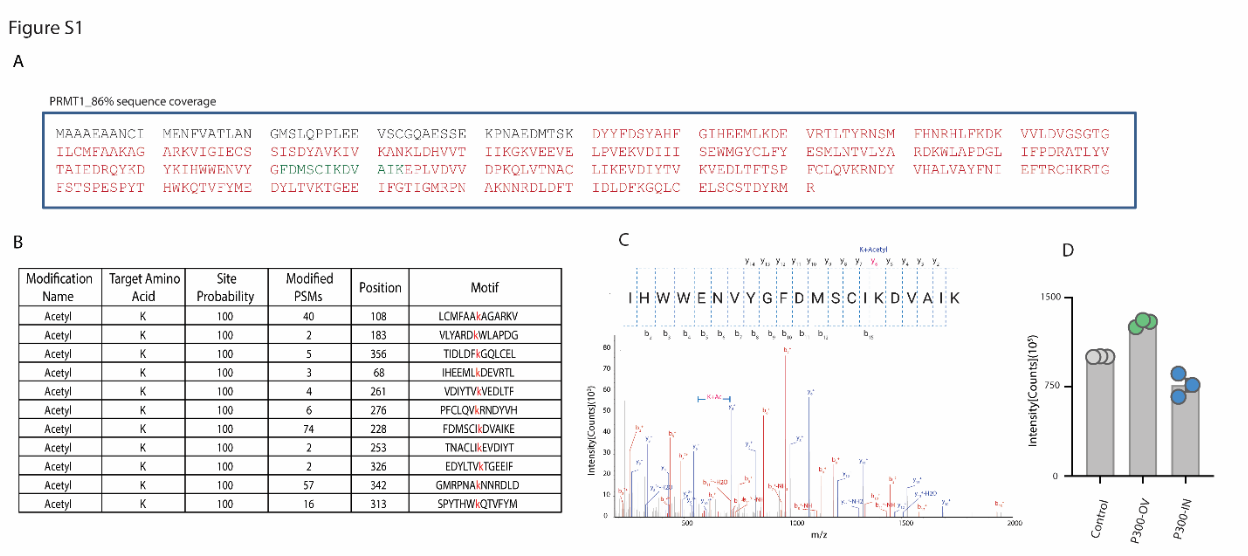


| Modification Name | Target Amino Acid | Site Probability | Modified PSMs | Position | Motif |
|---|---|---|---|---|---|
| Acetyl | K | 100 | 40 | 108 | LCMFAAkAGARKV |
| Acetyl | K | 100 | 2 | 183 | VLYARDkWLAPDG |
| Acetyl | K | 100 | 5 | 356 | TIDLDFkGQLCEL |
| Acetyl | K | 100 | 3 | 68 | IHEEMLkDEVRTL |
| Acetyl | K | 100 | 4 | 261 | VDIYTVkVEDLTF |
| Acetyl | K | 100 | 6 | 276 | PFCLQVkRNDYVH |
| Acetyl | K | 100 | 74 | 228 | FDMSCIkDVAIKE |
| Acetyl | K | 100 | 2 | 253 | TNACLIkEVDIYT |
| Acetyl | K | 100 | 2 | 326 | EDYLTVkTGEEIF |
| Acetyl | K | 100 | 57 | 342 | GMRPNAkNNRDLD |
| Acetyl | K | 100 | 16 | 313 | SPYTHWkQTVFYM |

**Figure S1. P300 acetylates PRMT1 at K228.** (A) Represents amino acid sequence of PRMT1 from IP-MS/MS experiments. Red indicates identified PRMT1 peptides and green indicates key peptide with acetyl K228. (B) Table shows the peptides identified with K-acetylation, peptides with two or more peptide-spectrum matches and 100 % site probability were included. (C) MS/MS spectra of peptide with acetyl K228. (D) Bar chart showing the intensity of peptide with acetyl K228 in control, p300 overexpressed and p300 inhibited condations. Label free quantification was used for quantification of peptides.

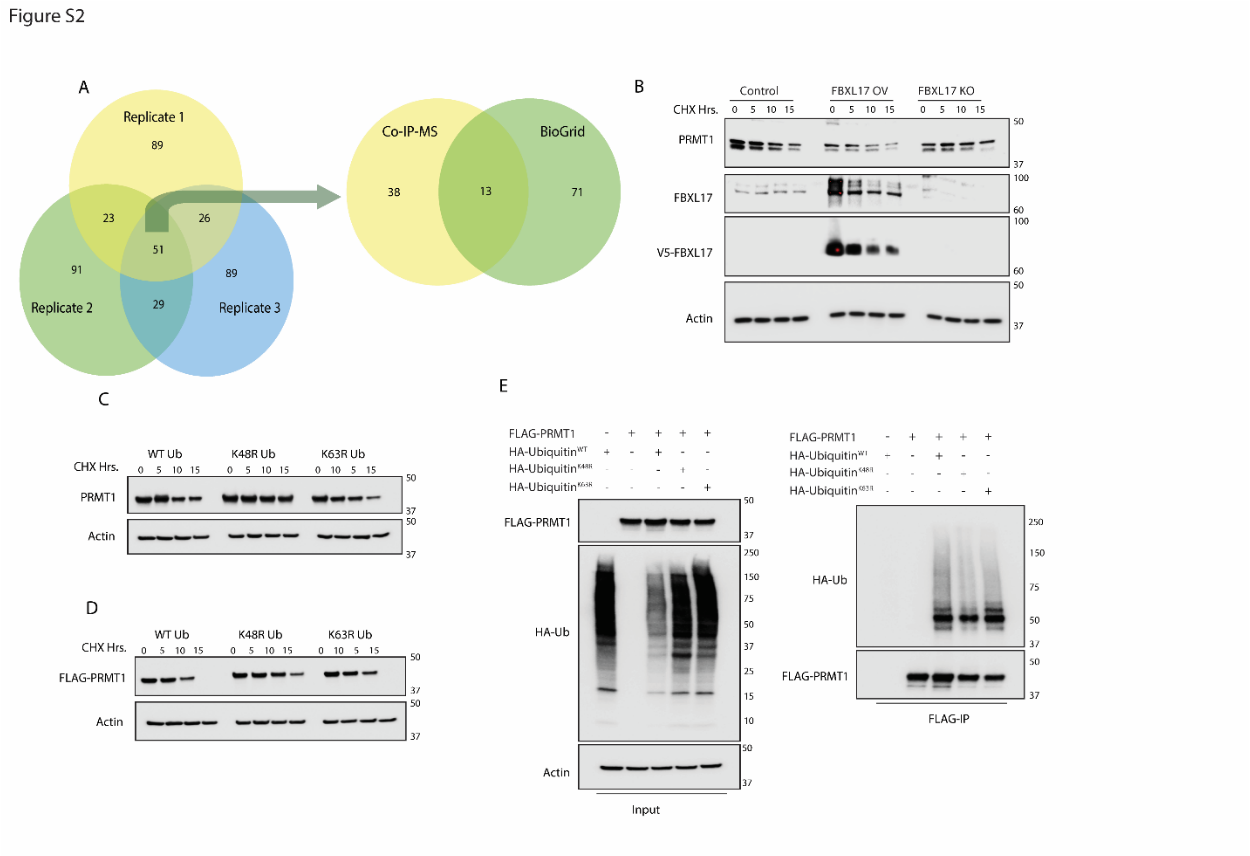


**Figure S2. FBXL17 decreases PRMT1 half-life by depositing K48 polyubiquitin chains.** (A) Venn diagram shows the interactors of FBXL17. Co-IP MS/MS of V5-FBXL17 was run in triplicates and later data was compared with the BioGrid database. (B) Western blotting shows the half-life of endogenous PRMT1 in control cells, FBXL17 overexpressed cells and FBXL17-KO cells under cycloheximide (50 µg/ml) chase conditions. (C) Western blotting shows the half-life of endogenous PRMT1 cells co-transfected with HA-Ubiquitin or HA-Ubiquitin mutants under cycloheximide (50 µg/ml) chase conditions. (C) Western blotting shows the half-life of overexpressed FLAG-PRMT1 cells co-transfected with HA-Ubiquitin or HA-Ubiquitin mutants under cycloheximide (50 µg/ml) chase conditions. (E) Western blotting shows the ubiquitination of FLAG-PRMT1 with co-transfection of HA-Ub WT and mutant K48R and K63R. FLAG-tagged PRMT1 was coexpressed with HA tagged WT, K48R and K63R ubiquitin (input), FLAG-tagged PRMT1 was immunoprecipitated using anti-FLAG affinity gel, followed by immunoblotting using anti-HA antibody to shows the ubiquitination status of PRMT1.

Figure S3

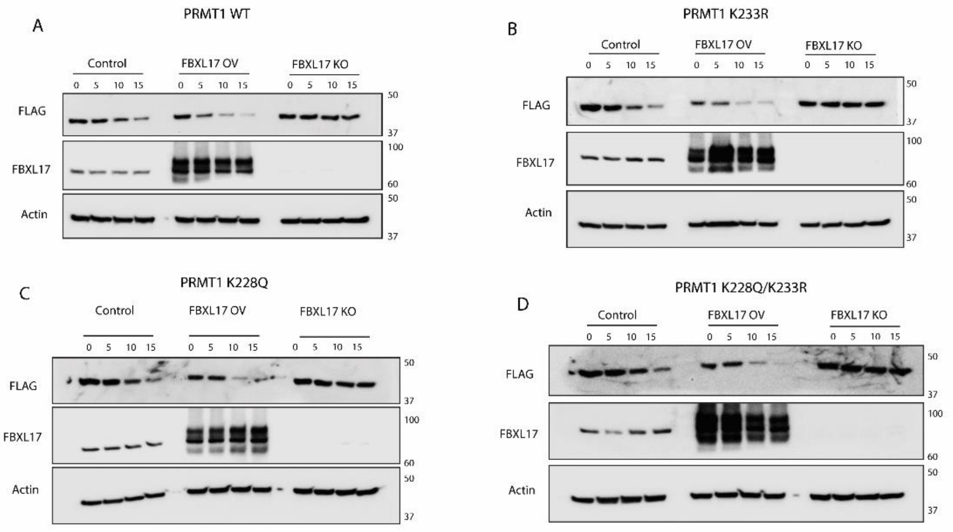


**Figure S3. PRMT1 acetylmimic mutations accelerate FBXL17-mediated degradation.** (A) Western blotting shows half-life of WT FLAG-tagged PRMT1 in FBXL17 overexpressed and KO conditions. (B) Western blotting shows half-life of K233R FLAG-tagged PRMT1 mutant in FBXL17 overexpressed and KO conditions. (C) Western blotting shows half-life of K228Q FLAG-tagged PRMT1 mutant in FBXL17 overexpressed and KO conditions. (D) Western blotting shows half-life of K228Q/K233R FLAG-tagged PRMT1 double mutant in FBXL17 overexpressed and KO conditions.

### List of Additional SI Files

| File | Contents |
|---|---|
| Table S1 | Protein List PRMT1 Co-IP |
| Table S2 | Protein List p300 Co-IP |
| Table S3 | PRMT1 Acetylation Sites |
| Table S4 | PRMT1 Ubiquitination Sites |
| Table S5 | Label Free Quantification of PRMT1 Acetylation Sites under p300 Overexpression/Inhibition Conditions |
| Table S6 | Protein List FBXL17 Co-IP |
| Table S7 | Label Free Quantification of PRMT1 GCE Peptides |
